# Validation of Fuze IMU system for ergonomics assessments

**DOI:** 10.1101/2022.12.05.519202

**Authors:** Elizabeth Serra-Hsu, Paolo Taboga

## Abstract

This study aims to validate the Fuze system (SwiftMotion, CA, USA), against the gold standard for motion capture, a 3D infra-red motion capture system (Vicon Nexus, Oxford, UK). Fuze system uses inertial measurement units and proprietary algorithms to calculate position and orientation of each body segment

Six subjects (3M and 3F) performed two activities that simulate common occupational physical activities. For both systems, we calculated the following joint angles: trunk relative to horizontal, left and right shoulder and hip joint angles. We also calculated the horizontal distance of each wrist relative to the the fifth lumbar vertebra. For each measurement, we calculated Bias (average difference between Fuze and Vicon system) and root mean squared error (RMSE). We also compared each measurement using a Statistical Parametric Mapping (SPM) method with a statistical significance level set at 0.05.

Compared to Vicon, Fuze system had a maximum Bias of 5.63 ± 1.60 degrees for the left shoulder angle and a maximum RMSE of 10.03 ± 2.73 degrees for the left hip angle. SPM analysis evidenced that for all the measurements, comparisons were within the critical thresholds for significance for the whole duration of the trials, indicating that we could not find a significant difference between Vicon and Fuze measurements.

In conclusion: the Fuze system compares well with the Vicon system and provides reliable data for the measurement of joint angles and body positions, that can be used in particular in non lab-based settings, for example in ergonomics risk assessments.

## Introduction

Movement studies have long since recognized motion capture systems as the gold standard of movement quantification, however they come with inherent limitations as the studies are restricted to laboratory settings (Poitras, et al. 2019). Development of portable, light weight sensors has been suggested as a solution to this problem. Inertial measurement units (IMUs), which consist of a variety of sensors including accelerometers, gyroscopes, and magnetometers that can measure kinetic and kinematic variables of motion, are being used more frequently for measurement of human motion. IMUs have many advantages as they may be covered with clothing, are easily transportable, and allow researchers to collect data from subjects in their natural environments. Through a number of validation studies, IMU kinematic data has been compared with motion capture data and deemed accurate through different methods, such as root mean square error (RMSE) (Mavor et al., 2020) and Bland-Altman limits of agreement (Robert-Lachaine et al., 2017). Simple movements for the upper and lower extremities, as well as axial skeleton have been validated (Poitras, et al. 2019). It has been shown that lower limb angles have a lower RMSE, between 2 and 11 degrees, while upper limb angles have a slightly higher RMSE, between 3 and 15 degrees (Mavor, et al. 2020, Robert-Lachaine, et al. 2017). While Bland-Altman biases have been shown to be between -1.10 deg and 1.35 deg with limits ranging from 0.96 deg to 5.06 deg (Teufl, et al. 2019).

The Fuze system (SwiftMotion, CA, USA), is a wearable device which quantifies the kinematics of the trunk and upper extremities using IMUs. This system contains eight IMUs that measure acceleration, velocity, and position of the trunk, arms, forearms, and thighs as well as two insoles that measure forces applied through each foot. Data from these sensors are transmitted to a tablet and used to summarize metrics that can be applied to risk assessment models commonly used in occupational health. These models can be applied to health and safety in the workplace with a strong focus on primary prevention of overuse injuries.

This study aims to validate the Fuze system against the gold standard for motion capture, a 3D infra-red motion capture system (Vicon Nexus, Oxford, UK). By comparing data collected by both systems, we can determine accuracy and reproducibility of the Fuze system, allowing researchers, occupational therapist, and investigators to select the most appropriate device for their future needs.

## Methods

### Participants

6 subjects, 3 male and 3 female, were recruited from within the Sac State student, staff, and faculty population. Subjects were between the ages of 18-55, with a heigh of 1.72 ± 0.11m, and body mass of 67.5 ± 16.0kg. They were in good health, and without reported musculoskeletal or neurological disease or disorder. Each subject gave written informed consent prior to participation. The protocol was approved by the University of California, Sacramento Institutional Review Board (IRB-19-20-88).

### Participant Preparation

Subjects were advised to wear their own comfortable clothes and shoes. The Fuze system, which consists of a lightweight harness consisting of Velcro straps, was then fitted over their clothing. The straps were fixed to the trunk, arms, hips, and thigh segments. IMUs were then attached to the harness for each body segment and the insoles were placed in each shoe. 10 IMUs were located in the following anatomical segments: trunk (at the level of second thoracic vertebra), pelvis (at the level of the fifth lumbar vertebra) bilateral upper arms (right above the elbow joints), bilateral lower arms (right above the wrist joints), bilateral thigh (mid-point between hip and knee joints), and bilateral lower leg (right above the ankle joints). Reflective markers were placed on each IMU as well as specific additional anatomical landmarks: clavicular notch, and bilaterally on acromion, lateral and medial elbow joints, lateral and medial wrist joints, anterior and posterior superior iliac spines, greater trochanter, lateral and medial knee joint, lateral and medial ankle joints (see Fig. 1).

**Fig. 1.**
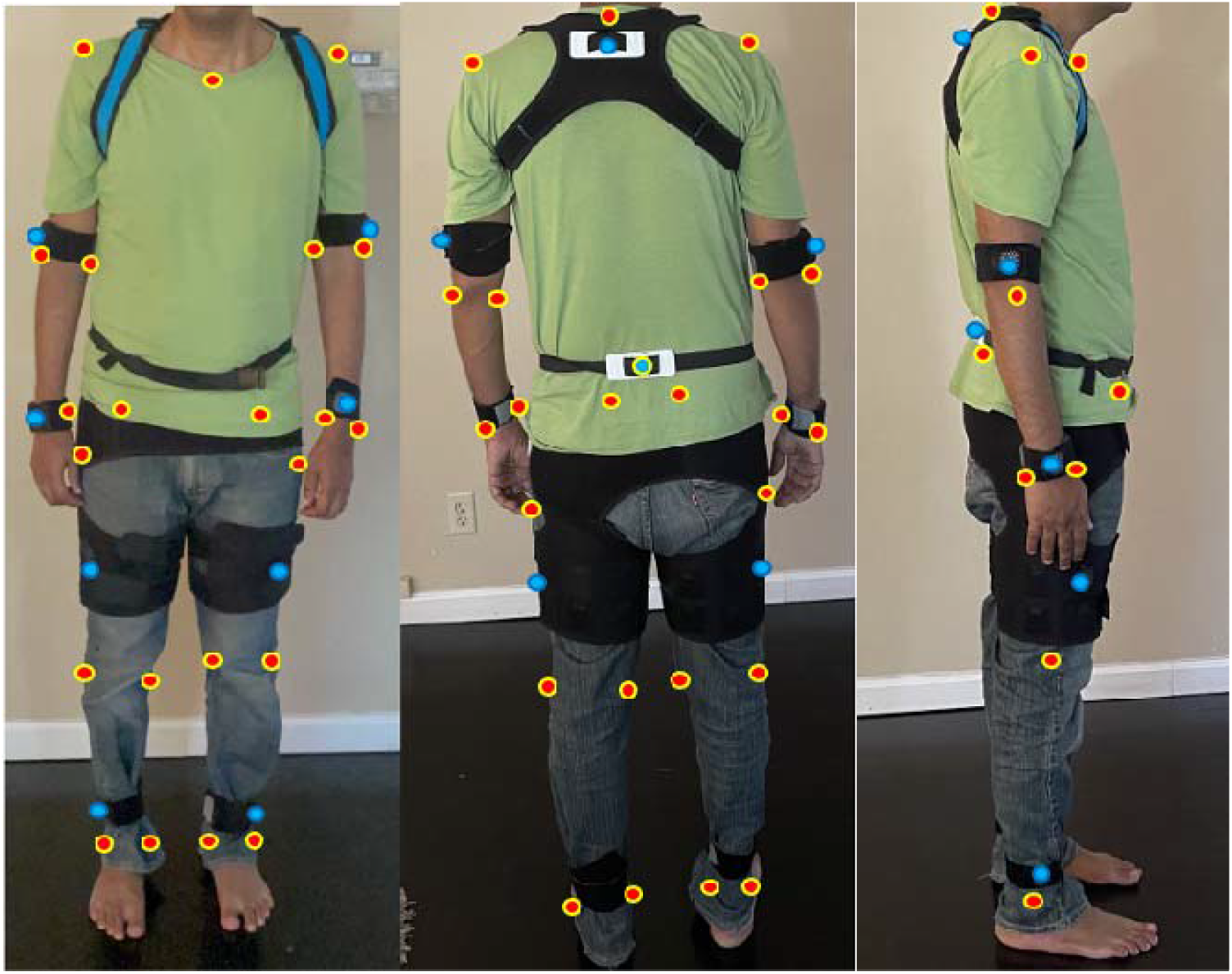
Subject wearing the harness with Velcro straps, the blue dots correspond to the 10 Fuze IMUs, the red dots correspond to the additional reflective markers used with the Vicon system.

### Movement Protocol

Each subject completed two activities that simulate common occupational physical activities. The first activity, labelled “static stoop”, consisted of a static hold of an empty box (0.4 m wide, 0.2 m deep and 0.2 m tall) with both hands at varying heights. The subject held the box at their shins, knees, and waist heights for 10 seconds each. The second activity, labelled “reaching”, consisted of standing at the edge of a table and leaning over to touch marks placed at 0.4 m, 0.6 m, 0.8 m and 1.0 m in front of the subject, and holding the position for 10 seconds at each mark.

### Data collection and analysis

The Fuze system (SwiftMotion, Berleley, USA) collected data at with a sampling rate of 100 Hz, the position and orientation of each IMU, and each relative anatomical segment, was calculated by means of a proprietary algorithm (SwiftMotion, Berleley, USA). The Vicon system (Vicon Nexus, Oxford, UK) consisted of 8 cameras uniformly distributed around the subject and mounted on the lab walls; we recorded the 3D position of each marker with the same sampling rate of 100 Hz. We then imported marker data in Visual 3D (C-Motion Inc., Germantown, MD) and, for each subject, we created anatomical models for the following segments: trunk, pelvis, left and right femur, left and right humerus, left and right forearm. For both systems, we then extracted the following angles in the sagittal plane: trunk relative to horizontal, shoulder joint angle as the angle of the humerus relative to the trunk, and hip joint angle as the angle of the femur relative to the pelvis. We also calculated the antero-posterior distance of each wrist IMU/marker relative to the IMU/marker placed on the fifth lumbar vertebra.

For both systems we filtered all data (markers and IMUs positions, segment and joint angles) with a Butterworth low-pass filter at 30 Hz. For each subject and for each movement the two different datasets, one from each system, were exported and synchronized by means of a custom Matlab algorithm.This synchronization allowed us to compare each data point at the same instant, irrespectively on when the data collection was started for each system.

After the datasets were synchronized, we extracted 100 points for each of the 10 seconds when the subject held the position, corresponding to a total of 300 data-points for the static stoop task (100 points x 3 heights) and 400 data-points for the reaching task (100 points x 4 positions). Each task was then time-normalized (0% representing the beginning of the movement, 100% representing the end).

Each variable (markers and IMUs positions, segments and joint angles) was then compared by calculating the following:

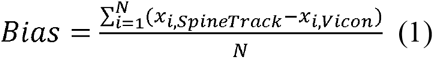

where *x_i,SpineTrack_* is the variable measured at the data-point number *i* by the Fuze system, *x_i,Vicon_* is the same variable measured by the Vicon system, and N is the total number of data points for each task. *Bias* represents therefore the average difference between Fuze and Vicon system; a positive value of *Bias*, for example, indicates that the Fuze system overestimates the Vicon measurement.

For each variable we also calculated the Root Mean Square Error (*RMSE*) between the two measurements as:

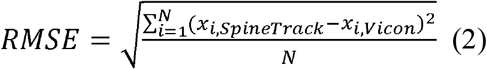

### Statistical analysis

We used a Statistical Parametric Mapping (SPM) method (Pataky et al., 2013) to compare trunk and bilateral hip angles for the static stoop task, and wrist positions, trunk and bilateral shoulder angles for the reaching tasks between Vicon and Fuze systems. Briefly, SPM methods allow the comparisons of two time-dependent variables for the whole dataset (with a number N of data-points collected at each time-normalized interval), this would be similar to performing a number of N t-tests for each data-point. SPM methods would therefore evidence if and when, e.g. at what time, the difference between the two variables would exceed a critical threshold calculated using the whole dataset, without the need to arbitrarily extract a global scalar representing the whole time-series (Pataky et al., 2013). We performed the SPM comparisons using a Matlab package (https://spm1d.org/), for each comparison statistical significance level was set at 0.05.

## Results

For each task, we report mean ± standard deviation of Bias and RMSE of each variable, calculated among all subjects (Table 1).

**Table 1.** Bias and RMSE of each analyzed variable for Reaching and Static stoop movements. For each variable, Bias represents the average difference between Fuze and Vicon systems (see Eq. 1) and RMSE is the root mean standard error between Fuze and Vicon systems (see Eq. 2), each variable is reported as mean ± s.d., calculated among all subjects. In **bold**, we highlight the maximum Bias and RMSE values among all variables.

| Movement | Variable | Bias | RMSE |
| --- | --- | --- | --- |
| Static stoop | Left Hip Angle (deg) | $4.21 \pm 4.81$ | **$10.03 \pm 2.73$** |
| | Right Hip Angle (deg) | $-0.73 \pm 4.81$ | $8.40 \pm 2.97$ |
| | Trunk Angle (deg) | $0.56 \pm 5.81$ | $7.50 \pm 3.51$ |
| Reaching | Left Shoulder Angle (deg) | **$5.63 \pm 1.60$** | $7.29 \pm 3.30$ |
| | Right Shoulder Angle (deg) | $-0.29 \pm 6.73$ | $6.82 \pm 4.33$ |
| | Trunk Angle (deg) | $2.76 \pm 3.35$ | $4.87 \pm 2.12$ |
| | Left Wrist Position (m) | $0.025 \pm 0.040$ | $0.052 \pm 0.024$ |
| | Right Wrist Position (m) | $0.029 \pm 0.049$ | $0.061 \pm 0.030$ |

For the static stoop task, we found that, compared to the Vicon system, the Fuze system overestimated left hip flexion angle by 4.21 ± 4.81 degrees, underestimated right hip flexion angle by -0.73 ± 4.81 and overestimated trunk angle by 0.56 ± 5.81 degrees.

For the reaching task, we found that compared to the Vicon system, the Fuze system overestimated the left shoulder angle by 5.63 ± 1.60 degrees, underestimated the right shoulder angle by -0.29 ± 6.73 degrees, overestimated the trunk angle by 2.76 ± 3.35 degrees, overestimated the left and right wrist positions by 0.025 ± 0.040 m and 0.029 ± 0.049 m respectively.

The SPM analysis evidenced that for the static stoop trials, left hip, right hip and trunk angles were all within the critical thresholds for significance for the whole duration of the trials, indicating that we could not find a significant difference between Vicon and Fuze measurements (Fig. 2).

**Figure 2.**
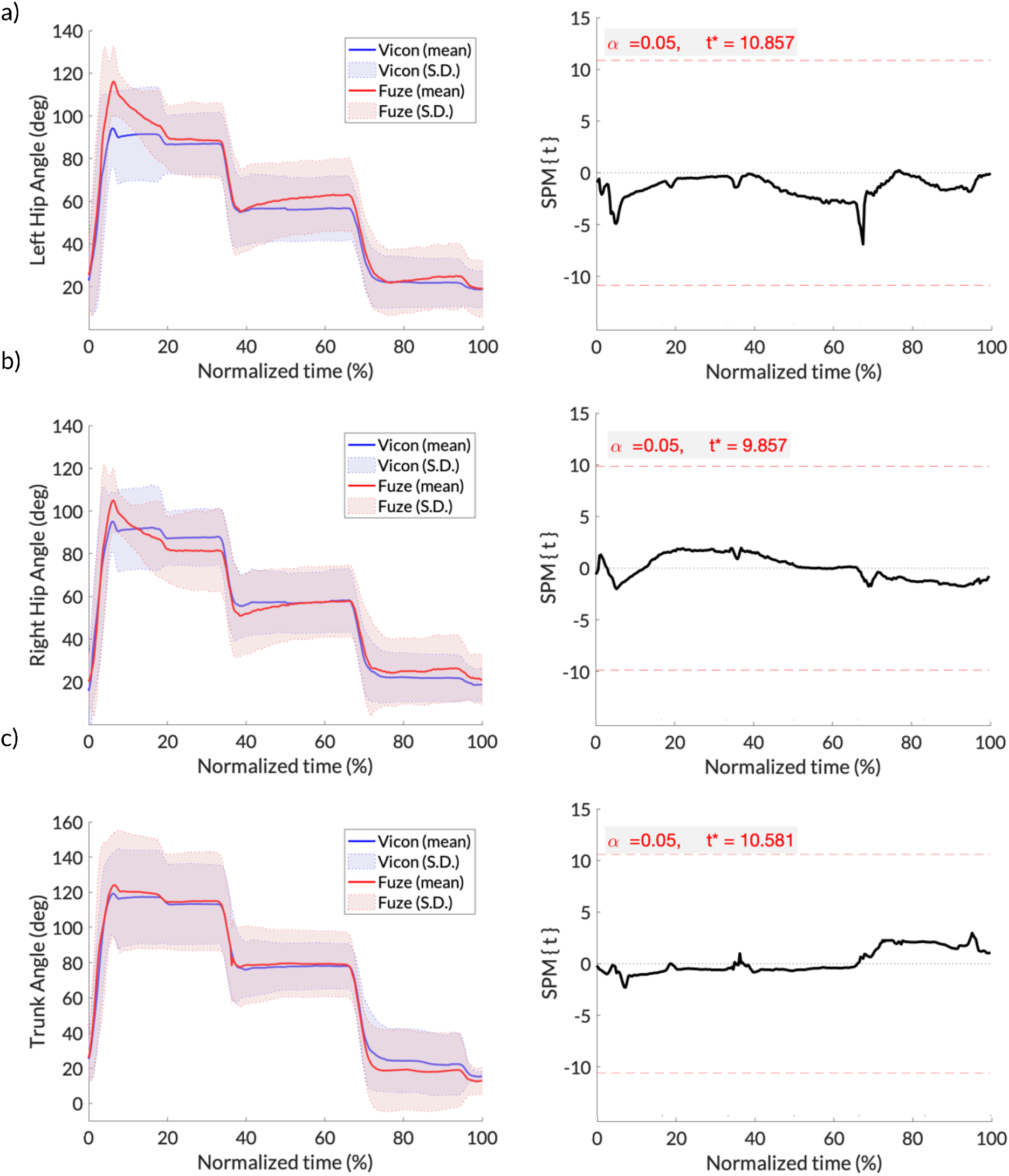
Static stoop trials. Left Hip (a), Right Hip (b) and Trunk (c), as a function of normalized time (0% beginning of trial, 100% end of trial). Left panels: the continuous lines represent the mean angle (in degrees), and the shaded areas represent the standard deviations of Vicon (blue) and Fuze (red) respectively. Right panels: the black lines represent the Statistical Parametric Mapping trajectories for each variable, the red dashed lines indicate the critical threshold for significance. All variables are within the thresholds for the whole duration of the trials.

Similarly, the SPM analysis evidenced that for the reaching trials, left shoulder, right shoulder and trunk angles, as well as left and right wrist positions were all within the critical thresholds for significance for the whole duration of the trials, indicating that we could not find a significant difference between Vicon and Fuze measurements (Fig. 3 and Fig. 4)

**Figure 3.**
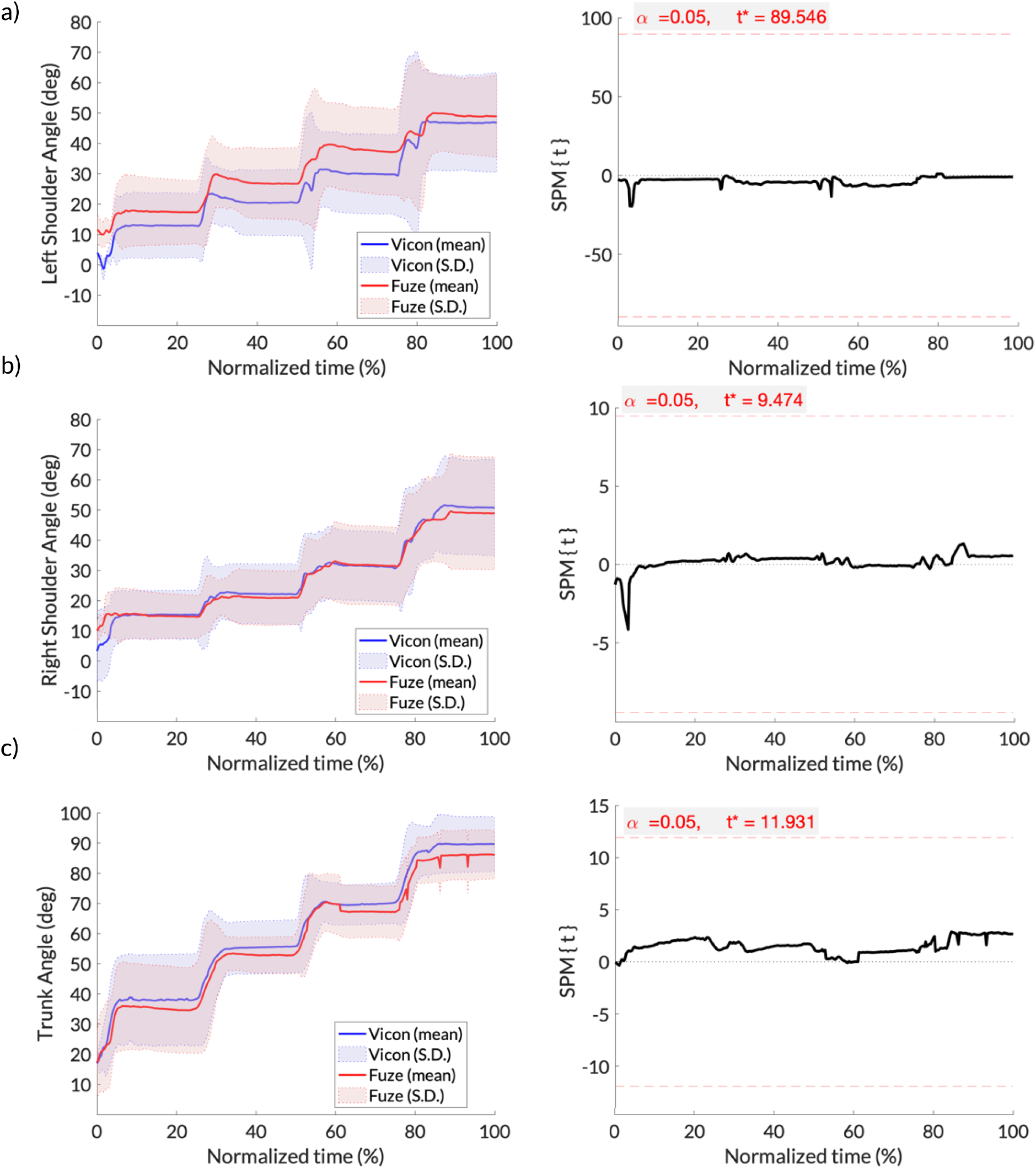
Reaching trials. Left Shoulder (a), Right Shoulder (b) and Trunk (c), as a function of normalized time (0% beginning of trial, 100% end of trial). Left panels: the continuous lines represent the mean angle (in degrees), and the shaded areas represent the standard deviations of Vicon (blue) and Fuze (red) respectively. Right panels: the black lines represent the Statistical Parametric Mapping trajectories for each variable, the red dashed lines indicate the critical threshold for significance. All variables are within the thresholds for the whole duration of the trials.

**Figure 4.**
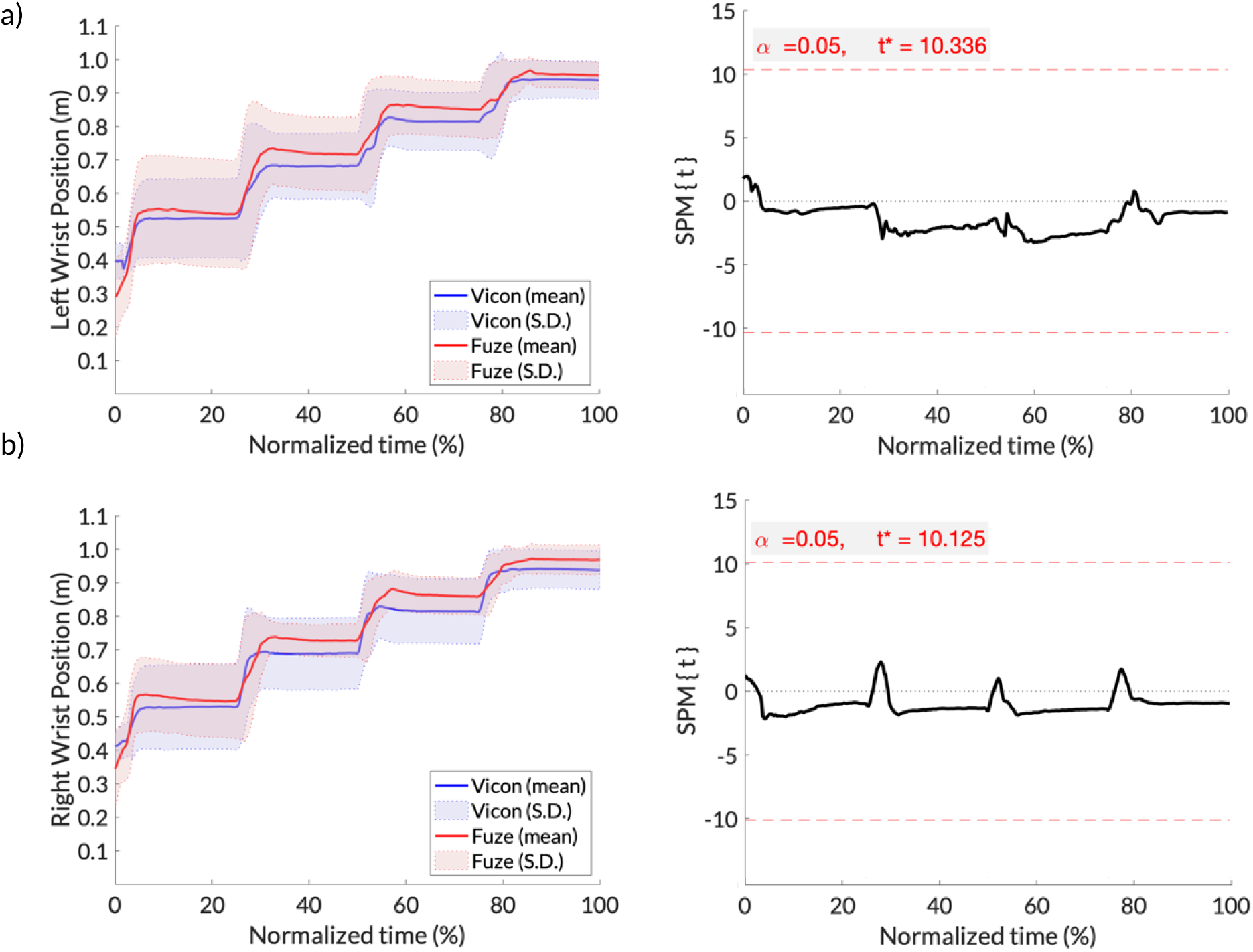
Reaching trials. Left Wrist (a), Right Wrist (b), as a function of normalized time (0% beginning of trial, 100% end of trial). Left panels: the continuous lines represent the mean position (in meters), and the shaded areas represent the standard deviations of Vicon (blue) and Fuze (red) respectively. Right panels: the black lines represent the Statistical Parametric Mapping trajectories for each variable, the red dashed lines indicate the critical threshold for significance. All variables are within the thresholds for the whole duration of the trials.

## Discussion and conclusions

The maximum Bias, calculated as the mean difference between the Fuze and Vicon systems, corresponded to the left shoulder flexion angle in the reaching trials (Bias = 5.63 ± 1.60 degrees). As seen in Figure 3, panel a, the Fuze system appears to overestimate the shoulder flexion for almost all data points, as opposed to the right shoulder angle where the lines for each system appear to overlap for the majority of the data points (Bias = -0.29 ± 6.73 degrees). Given that the only difference between right and left IMUs is their positioning on the subject, and not how the two shoulder flexion angles are measured/calculated, we could explain this difference as slightly misplaced IMU sensor on one or more subject on the left upper arm. This difference highlights the importance of accurate placement of each IMU sensor on each body segment: given that only one sensor is needed to calculate the position and orientation of that segment, a misplaced/not secured sensor could lead to inaccurate data.

The maximum RMSE values corresponded to the left hip flexion angle (10.03 ± 2.73 degrees) and right hip flexion angle (8.40 ± 2.97 degrees), in the static stoop trials. This is in accord with previous studies: (Mavor et al., 2020) for example reported RMSE values of 8.57 ± 3.89 and 8.43 ± 4.29 degrees for the left and right hip flexion angles respectively.

Overall, whether the agreement between the Fuze and Vicon systems is acceptable or not depends on the application. An error of less than 6 degrees when evaluating shoulder flexion angles is negligible, in particular in ergonomics risk assessments. For example the RULA (rapid upper limb assessment, (McAtamney & Nigel Corlett, 1993)) scores shoulder positions based on comparatively wide ranges (1: 0-20 degrees, 2: 20-45 degrees, 3: 45-90 degrees or 4: more than 90 degrees). Similarly, the OCRA assessment (Occhipinti, 1998) evaluates shoulder positions based on broad flexion ranges (20, 40, 60 or more than 60 degrees). Similarly, an error (Bias) of less than 0.03 m in the position of the wrist (Table 1) would be negligible when calculating the NIOSH lift index (Waters et al., 2007).

We must point out that while it is generally assumed that optical systems are the ‘gold standard’ in motion capture, they are not error-free: marker misplacement, limited field of view of the cameras, limited capture volume etc. are only a few of the inherent limitations of these systems. The use of IMU-based systems allows to overcome, at least partially, these limitations: IMUs don’t need to be in direct line-of-sight with a receiver, they can be covered with clothes or Velcro straps that secure the to each body part, allow for faster data processing etc., all features that are desired in “real world” situations such as workplace ergonomics evaluations, outdoors etc.

In conclusion the Fuze system compares well with the Vicon system and provides reliable data for the measurement of joint angles and body positions, that can be used in particular in non lab-based settings where the performance of other systems, such as optical systems, would be inferior.

## REFERENCES

Mavor, M. P., Ross, G. B., Clouthier, A. L., Karakolis, T., & Graham, R. B. (2020). Validation of an IMU Suit for Military-Based Tasks. Sensors (Basel), 20(15). https://doi.org/10.3390/s20154280

McAtamney, L., & Nigel Corlett, E. (1993). RULA: a survey method for the investigation of work-related upper limb disorders. Appl Ergon, 24(2), 91–99. https://doi.org/10.1016/0003-6870(93)90080-s

Occhipinti, E. (1998). OCRA: a concise index for the assessment of exposure to repetitive movements of the upper limbs. Ergonomics, 41(9), 1290–1311. https://doi.org/10.1080/001401398186315

Pataky, T. C., Robinson, M. A., & Vanrenterghem, J. (2013). Vector field statistical analysis of kinematic and force trajectories. J Biomech, 46(14), 2394–2401. https://doi.org/10.1016/j.jbiomech.2013.07.031

Robert-Lachaine, X., Mecheri, H., Larue, C., & Plamondon, A. (2017). Validation of inertial measurement units with an optoelectronic system for whole-body motion analysis. Med Biol Eng Comput, 55(4), 609–619. https://doi.org/10.1007/s11517-016-1537-2

Waters, T. R., Lu, M. L., & Occhipinti, E. (2007). New procedure for assessing sequential manual lifting jobs using the revised NIOSH lifting equation. Ergonomics, 50(11), 1761–1770. https://doi.org/10.1080/00140130701674364

